# Matrin 3-dependent neurotoxicity is modified by nucleic acid binding and nucleocytoplasmic localization

**DOI:** 10.1101/268458

**Authors:** Ahmed M. Malik, Roberto A. Miguez, Xingli Li, Ye-Shih Ho, Eva L. Feldman, Sami J. Barmada

## Abstract

Abnormalities in nucleic acid processing are associated with the development of amyotrophic lateral sclerosis (ALS) and frontotemporal dementia (FTD). Mutations in *Matrin 3* (*MATR3*), a poorly understood DNA- and RNA-binding protein, cause familial ALS/FTD, and MATR3 pathology is a feature of sporadic disease, suggesting that MATR3 dysfunction is integrally linked to ALS pathogenesis. Using a primary neuron model to assess MATR3-mediated toxicity, we noted that neurons were bidirectionally vulnerable to MATR3 levels, with pathogenic MATR3 mutants displaying enhanced toxicity. MATR3’s zinc finger domains partially modulated toxicity, but elimination of its RNA recognition motifs had no effect on neuronal survival, instead facilitating its self-assembly into liquid-like droplets. In contrast to other RNA-binding proteins associated with ALS, cytoplasmic MATR3 redistribution mitigated neurodegeneration, suggesting that nuclear MATR3 mediates toxicity. Our findings offer a foundation for understanding MATR3-related neurodegeneration and how nucleic acid binding functions, localization, and pathogenic mutations drive sporadic and familial disease.

## INTRODUCTION

Amyotrophic lateral sclerosis (ALS) is a progressive neurodegenerative disorder resulting in the death of upper and lower motor neurons (Charcot and Joffroy, 1869). Mounting evidence indicates that RNA-binding proteins (RBPs) are integrally involved in the pathogenesis of ALS (Taylor et al., 2016). The majority (>95%) of ALS patients display cytoplasmic mislocalization and deposition of the RBP TDP-43 (TAR DNA/RNA-binding protein of 43 kDa) in affected tissue (Neumann et al., 2006). Moreover, over 40 different ALS-associated mutations have been identified in the gene encoding TDP-43, and mutations in several different RBPs have been similarly linked to familial ALS (Kabashi et al., 2008; Kwiatkowski et al., 2009; Vance et al., 2009; Barmada and Finkbeiner, 2010; Ticozzi et al., 2011; Kim et al., 2013). These mutations often cluster in intrinsically disordered domains that facilitate reversible liquid-liquid phase separation (LLPS), thereby creating ribonucleoprotein granules important for RNA processing, shuttling of mRNAs to sites of local translation, or sequestration of transcripts during stress. Pathogenic mutations in the genes encoding TDP-43 and related RBPs, including FUS and TIA1, shift the equilibrium towards irreversible phase separation and the formation of cytoplasmic aggregates analogous to those observed in post-mortem tissues from patients with ALS (Johnson et al., 2009; Patel et al., 2015; Gopal et al., 2017; Mackenzie et al., 2017). The downstream implications of abnormal LLPS on RNA misprocessing, RBP pathology, and neurodegeneration in ALS are unknown, however.

Matrin 3 (MATR3) is a DNA- and RNA-binding protein with wide-ranging functions in nucleic acid metabolism including gene transcription, the DNA damage response, splicing, RNA degradation, and the sequestration of hyperedited RNAs (Belgrader et al., 1991; Hibino et al., 2000; Zhang and Carmichael, 2001; Salton et al., 2014; Coelho et al., 2015; Rajgor et al., 2016; Uemura et al., 2017). The MATR3 S85C mutation leads to autosomal dominant distal myopathy with vocal cord and pharyngeal weakness (Feit et al., 1998; Senderek et al., 2009). A more recent report reclassified a subset of patients with this diagnosis as having ALS and noted several additional MATR3 mutations in individuals with ALS and frontotemporal dementia (FTD), placing MATR3 in a group of proteins implicated in familial ALS, FTD, and myopathy; other members of this family include VCP, TIA1 and hnRNPA2/B1 (Kimonis et al., 2008; Johnson et al., 2010; Kim et al., 2013; Klar et al., 2013; Johnson et al., 2014; Mackenzie et al., 2017). A total of 13 pathogenic MATR3 mutations have now been identified, most of which are located in disordered stretches of the protein (Fig. 1A) (Millecamps et al., 2014; Origone et al., 2015; Leblond et al., 2016; Xu et al., 2016; Marangi et al., 2017). Additionally, post-mortem analyses demonstrated MATR3 pathology—consisting of cytoplasmic MATR3 accumulation as well as strong nuclear immunostaining—in patients with sporadic ALS and familial disease due to *C9orf72* hexanucleotide expansions and *FUS* mutations (Dreser et al., 2017; Tada et al., 2017).

**Figure 1.**
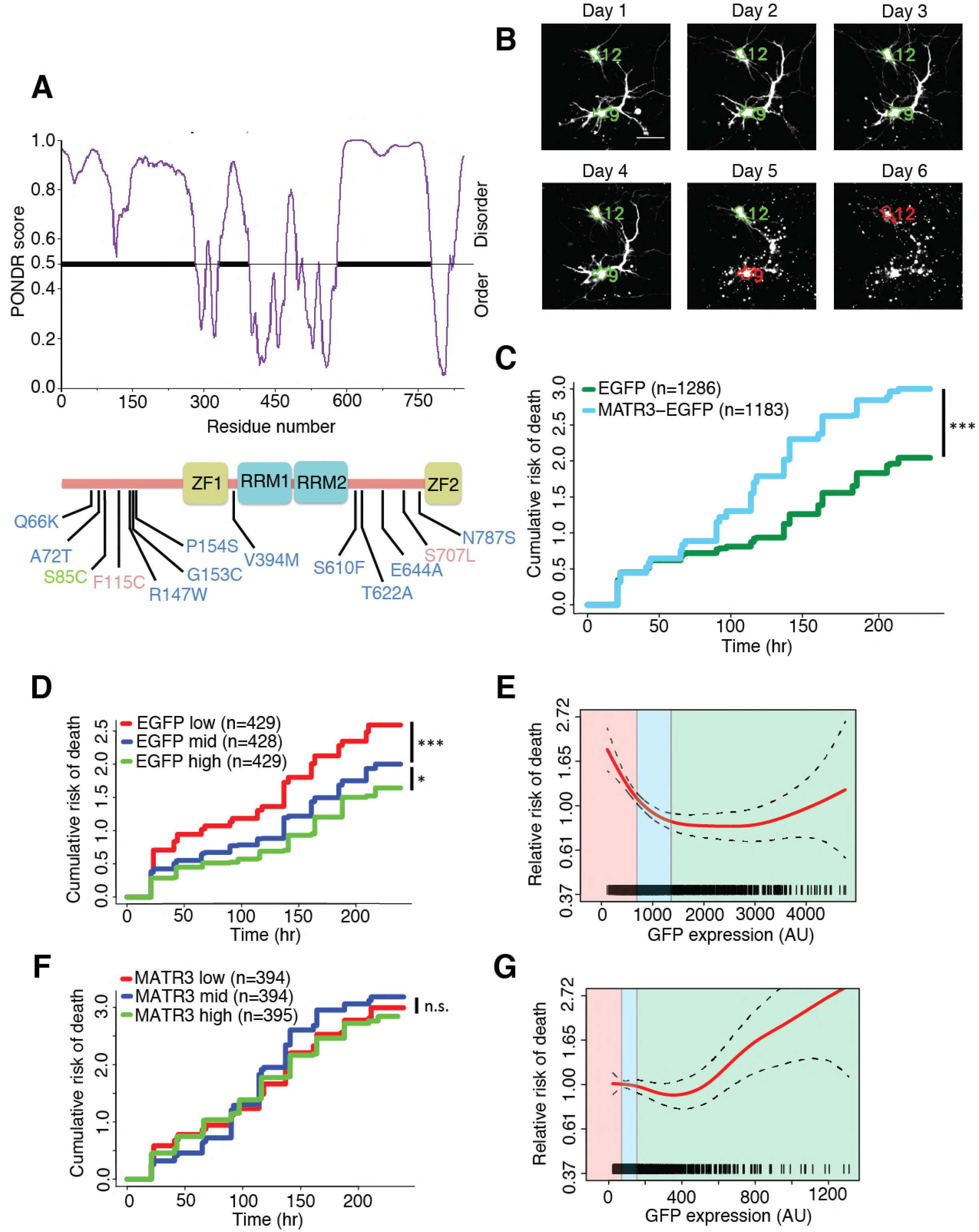
MATR3 overexpression results in dose-dependent neurodegeneration. **A.** Diagram of MATR3 showing nucleic acid-binding domains as well as the distribution of pathogenic mutations implicated in ALS (blue), ALS/FTD (red), and ALS/distal myopathy (green) within domains predicted to be disordered by PONDR VSL2 (Peng et al., 2006). **B.** Longitudinal fluorescence microscopy (LFM) allows unique identification and tracking of thousands of primary neurons (green outlines) transfected with fluorescent proteins, as well as monitoring of cell death (red outlines), indicated by loss of fluorescence signal and changes in morphology. Scale bar, 20 μm. **C.** MATR3-EGFP expressing neurons exhibited a higher risk of death compared to neurons expressing only EGFP, as quantified by the hazard ratio (HR) (HR = 1.48, EGFP n = 1286, MATR3-EGFP n = 1183; p < 2 × 10^−16^; Cox proportional hazards). **D.** EGFP expressing cells were divided into three equal groups based off expression level. Increased survival was associated with higher expression levels of EGFP (comparing to medium expressers n = 428: low expressers HR = 1.39, n = 429, p = 4.2 × 10^−5^; high expressers HR = 0.79, n = 429, p = 0.024; Cox proportional hazards). **E.** Penalized spline modeling confirmed a protective effect associated with higher EGFP expression that plateaus at ~1500 arbitrary units (AU); shaded colors represent low, medium and high expression ranges as in (**D**) (p = 5.3 × 10^−6^; penalized spline regression). **F.** There were no significant differences in survival among neurons expressing low, medium, or high levels of MATR3-EGFP (comparing to medium expressers n = 394: low expressers HR = 0.99, n = 394, p = 0.94; high expressers HR = 1.06, n = 395, p = 0.54; Cox proportional hazards). **G.** Similarly, penalized spline analysis showed no relationship between expression and survival at low and medium expression but a significant increase in risk of death with high MATR3-EGFP levels (p = 0.012; penalized spline regression).

Together, these observations suggest that MATR3 may be a common mediator of disease even in those without *MATR3* mutations. Even so, little is known about MATR3’s functions in health or in disease, and the mechanisms underlying MATR3-dependent neurotoxicity remain unclear. Here, we establish an *in vitro* model of MATR3-mediated neurodegeneration and take advantage of this model to investigate the intrinsic properties and domains of MATR3 required for toxicity. Furthermore, we examine how disease-associated MATR3 mutations affect these properties to enhance neurodegeneration.

## RESULTS

### MATR3 levels modulate neuronal survival in an *in vitro* model of neurodegeneration

We first asked how MATR3 expression is related to neurodegeneration using longitudinal fluorescence microscopy (LFM), a sensitive high-content imaging system that we assembled for assessing neuronal function and survival at the single-cell level. As *MATR3* mutations cause a spectrum of disease that includes ALS and FTD, we modeled neurotoxicity in primary mixed cortical cultures, a system that recapitulates key features of ALS/FTD pathogenesis (Barmada et al., 2010; Barmada et al., 2014; Barmada et al., 2015). Primary neurons were transfected with diffusely localized mApple to enable visualization of neuronal cell bodies and processes by fluorescence microscopy. In addition, cells were co-transfected with constructs encoding enhanced green fluorescent protein (EGFP) or MATR3 fused with EGFP. Cultures were imaged by fluorescence microscopy at 24 h intervals for 10 days, and custom scripts used to generate uniquely labeled regions of interest (ROIs) corresponding to each cell (Fig. 1B). Rounding of the soma, retraction of neurites or loss of fluorescence indicated cell death; these criteria proved to be sensitive markers of neurodegeneration in previous studies (Arrasate and Finkbeiner, 2005). We used the time of death for individual cells to calculate an overall risk of death, expressed as a hazard ratio (HR), corresponding to the likelihood of cell death in each population relative to a control or reference group (Christensen, 1987). In doing so, we observed that MATR3(WT)-EGFP overexpression significantly increases the risk of death compared to EGFP alone, with a HR of 1.48 (Fig. 1C).

Next, we investigated the dose-dependency of this MATR3 toxicity through two alternative but complementary approaches. Transient transfection delivers a different amount of vector to each cell, resulting in substantial variability in protein expression for individual cells. Since fluorescence intensity is directly proportional to fluorophore levels (Arrasate et al., 2004), the GFP intensity within each ROI provides an estimate of EGFP or MATR3(WT)-EGFP expression for individual neurons. Based on the GFP intensity measured 24 h after transfection, we divided transfected neurons into three groups: those that expressed low, medium, and high levels of EGFP or MATR3(WT)-EGFP. We then assessed the relative survival of these groups over time, and compared the risk of death in each by Cox proportional hazards. In doing so, we noted that cells that express low EGFP levels display an increased risk of death compared to those in the medium or high EGFP expression categories, potentially due to poor protein expression by unhealthy or dying cells (Fig. 1D). We also analyzed the relationship between GFP intensity and survival using penalized splines, which approximate both linear and non-linear relationships by treating GFP intensity as a continuous variable (Miller et al., 2010; Barmada et al., 2015). In this model, increasing EGFP expression predicted improved survival, but the effect plateaued at approximately 1500 arbitrary units (AU) (Fig. 1E). These data imply that lower expression of a neutral protein such as EGFP is tied to reduced survival, consistent with the results of previous studies (Miller et al., 2010; Barmada et al., 2015).

To determine how MATR3(WT)-EGFP expression is related to neuronal survival, we likewise separated neurons into three groups (low, medium and high) depending on MATR3(WT)-EGFP levels and assessed survival in each group. Unlike cells expressing EGFP alone, we detected no significant difference in survival between the low, medium, and high MATR3(WT)-EGFP expression groups (Fig. 1F). Correspondingly, the penalized spline model shows no clear relationship between risk of death and MATR3(WT)-EGFP levels for cells displaying low or medium GFP intensity. However, in contrast to cells expressing EGFP alone, we noted an increase in the risk of death with high MATR3(WT)-EGFP expression (Fig. 1G), suggesting that the extended survival observed in high-expressing cells is offset by the production of a toxic protein. Taken together, these data support a dose-dependent toxicity of MATR3(WT) in primary neurons.

Several *MATR3* mutations have been associated with familial ALS, FTD, and hereditary distal myopathy (Senderek et al., 2009; Johnson et al., 2014; Millecamps et al., 2014; Origone et al., 2015; Leblond et al., 2016; Xu et al., 2016; Marangi et al., 2017). To determine if disease-associated *MATR3* mutations accentuate neurodegeneration, we created MATR3-EGFP fusion proteins harboring one of four mutations originally implicated in familial disease: S85C, F115C, P154S, and T622A (Fig. 1A). Primary rodent cortical neurons expressing these mutant MATR3-EGFP constructs exhibited the same granular nuclear distribution as MATR3(WT)-EGFP, without obvious aggregation or cytoplasmic mislocalization, consistent with prior reports (Fig. 2A) (Gallego-Iradi et al., 2015; Boehringer et al., 2017). Even so, all four displayed a subtle but significant increase in toxicity over MATR3(WT)-EGFP when overexpressed in primary neurons (Fig. 2B), consistent with either gain-of-function or dominant negative loss-of-function mechanisms contributing to mutant MATR3-associated neurodegeneration.

**Figure 2.**
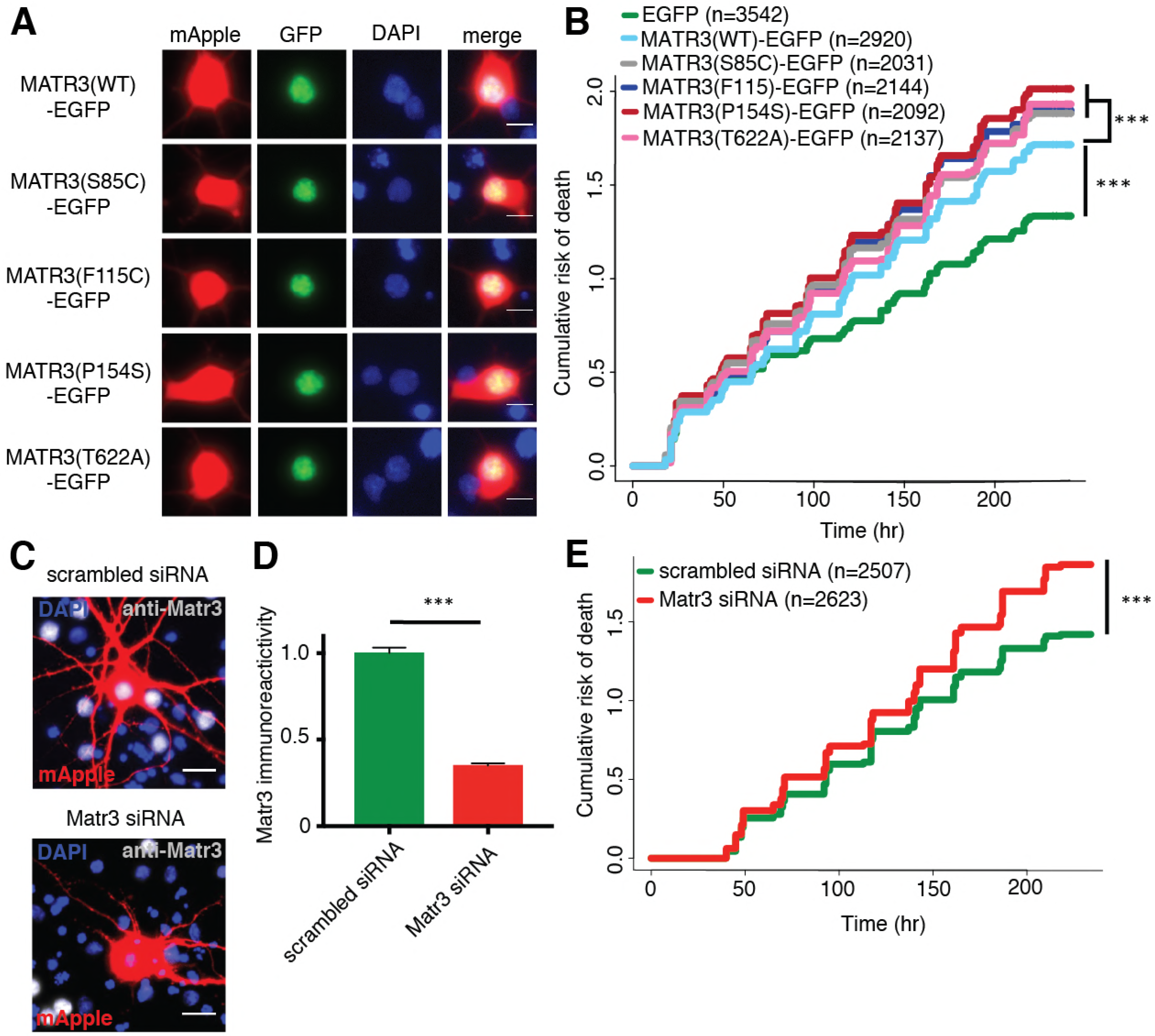
Neurons are susceptible to both gain-of-function and loss-of-function MATR3 toxicity. **A.** In primary rodent cortical neurons, the S85C, F115, P154S, and T622A disease-associated MATR3 mutants have the same granular nuclear distribution as MATR3(WT)-EGFP. **B.** All four disease mutations display a subtle but significant increase in toxicity compared to MATR3(WT)-EGFP (comparing to MATR3(WT)-EGFP n = 2920; MATR3(S85C)-EGFP HR = 1.16, n = 2031, p = 3.79 × 10^−6^; MATR3(F115C)-EGFP HR = 1.14, n = 2144, p = 5.57 × 10^−5^; MATR3(P154S)-EGFP HR = 1.24, n = 2092, p = 1.77 × 10^−11^; MATR3(T622A)-EGFP HR = 1.14, n = 2137, p = 6.02 × 10^−5^; Cox proportional hazards). **C-D.** siRNA targeting the endogenous rat *Matr3* reduced MATR3 antibody reactivity by approximately 65% (scrambled siRNA n = 576, anti-Matr3 siRNA n = 508, p < 0.0001; two-tailed t-test). **E.** Neurons transfected with anti-Matr3 siRNA displayed a higher risk of death compared to those transfected with scrambled siRNA (HR = 1.20, scrambled siRNA n = 2507, anti-Matr3 n = 2623, p = 2.05 × 10^−8^; Cox proportional hazards). Scale bars in (**A**), 10 μm; scale bars in (**C**), 20 μm.

To determine if loss of endogenous MATR3 function is sufficient for neurodegeneration, we transfected primary neurons with mApple and siRNA targeting the amino (N)-terminal coding region of rodent *Matr3* or a scrambled siRNA control. Three days after transfection, Matr3 immunoreactivity was used to quantify efficacy of knockdown in transfected cells (Fig. 2C). Compared to scrambled siRNA-transfected cells, we noted consistent depletion of the endogenous rat Matr3 by approximately 65% in those transfected with siRNA targeting *Matr3* (Fig. 2D). Having confirmed knockdown, we imaged a separate set of transfected cells for 10 days to assess the effect of *Matr3* knockdown on neuronal survival. In doing so, we observed a 20% increase in the risk of death upon *Matr3* depletion in comparison to scrambled siRNA (Fig. 2E). These data suggest that neurons are vulnerable to both increases and decreases in MATR3 levels and function; further, pathogenic *MATR3* mutations may elicit neurodegeneration via gain- or loss-of-function mechanisms, or through elements of both.

### MATR3’s zinc finger domains modulate overexpression toxicity, but its RNA recognition motifs mediate self-association

To identify the functional domains involved in MATR3-mediated neurodegeneration, we systematically deleted each of the annotated MATR3 domains and evaluated subsequent toxicity upon overexpression in primary neurons (Fig. 3A). MATR3 has two zinc-finger (ZF) domains of the C2H2 variety, which bind DNA but may also recognize RNA and/or mediate protein-protein interactions (Brayer et al., 2008; Burdach et al., 2012). Deletions of ZF1, ZF2, or both had no observable effect on MATR3-EGFP localization (Fig. 3B), and ZF1 deletion by itself did not significantly alter toxicity compared to full-length MATR3-EGFP. In contrast, ZF2 deletion, either in isolation or combined with ZF1 deletion, partially rescued MATR3-EGFP overexpression toxicity (Fig. 3C).

**Figure 3.**
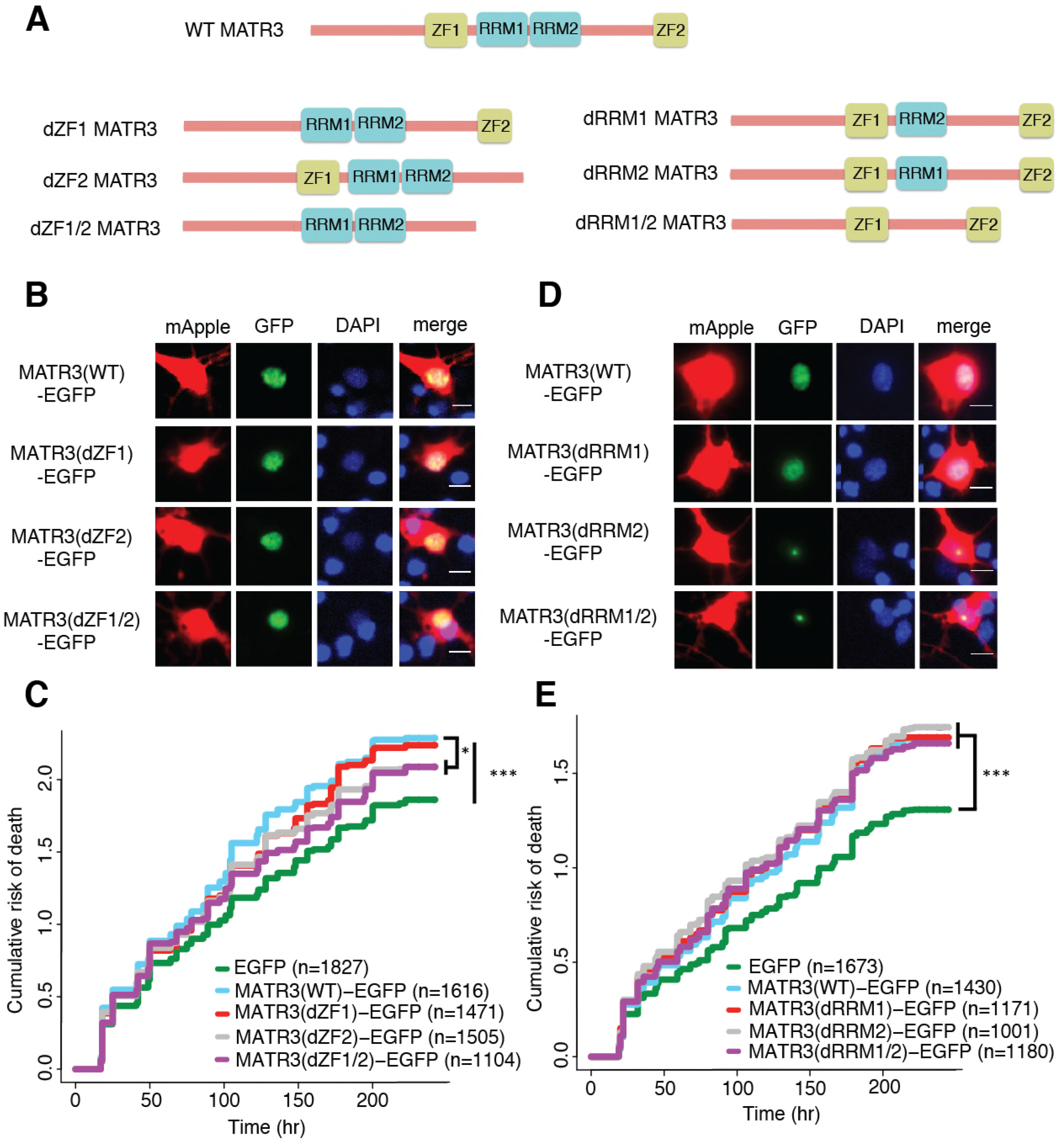
MATR3’s ZFs mediate overexpression toxicity, and its RRMs regulate subcellular distribution. **A.** Schematic of MATR3 domain deletion mutants. **B.** Zinc finger (ZF) domain deletions do not change the localization of MATR3-EGFP compared to the full-length protein. **C.** ZF2 deletion, either in isolation or combination with ZF1, results in modest rescue of overexpression toxicity (comparing to MATR3(WT)-EGFP n = 1616: MATR3(dZF1)-EGFP HR = 0.94, n = 1471, p = 0.10; MATR3(dZF2)-EGFP HR =0.93, n = 1505, p = 0.040; MATR3(dZF1/2)-EGFP HR = 0.90, n = 1104, p = 0.0093; Cox proportional hazards). **D.** While MATR3(dRRM1)-EGFP exhibits the same localization as MATR3(WT)-EGFP, deletion of RRM2 results in redistribution into intranuclear granules. **E.** RRM deletion had little effect on MATR3-mediated toxicity (comparing to MATR3(WT)-EGFP n = 1430: MATR3(dRRM1)-EGFP HR = 1.05, n = 1171, p = 0.25; MATR3(dRRM2)-EGFP HR = 1.09, n = 1001, p = 0.066; MATR3(dRRM1/2)-EGFP HR = 1.04, n = 1180, p = 0.42). Scale bars in (**B**) and (**D**), 10 μm.

We next created deletion variants of MATR3’s RNA recognition motifs (RRMs) to test their contribution to MATR3-mediated neurodegeneration. As with the MATR3 ZF domains, RRMs are capable of recognizing both RNA and DNA (Inagaki et al., 1996). While deletion of RRM1 failed to affect MATR3-EGFP localization, we noted a striking redistribution of MATR3(dRRM2)-EGFP into intranuclear granules in a subset of transfected neurons (Fig. 3D). Deletion of RRM1 in combination with RRM2 produced the same phenotype, suggesting that RRM2 normally prevents such redistribution. These nuclear granules formed by MATR3(dRRM2)-EGFP and MATR3(dRRM1/2)-EGFP were uniformly spherical in shape, and their presence was accompanied by a reduction in the intensity of diffusely-distributed MATR3 within the nucleus, suggesting that they represent hyperconcentrated MATR3 puncta. Evidence from previous studies indicates that RNA recognition by MATR3 may be largely—but not solely—driven by RRM2 (Hibino et al., 2006; Salton et al., 2011). Consistent with this, our finding that RRM2 deletion induces the formation of nuclear condensates suggests that RNA binding normally keeps MATR3 diffuse by preventing an intrinsic tendency for self-association. Despite the dramatic shift in MATR3-EGFP distribution with RRM2 deletion, there was no associated change in the toxicity of MATR3-EGFP lacking RRM1, RRM2 or both in comparison to MATR3(WT)-EGFP (Fig. 3E). This finding stands in contrast to what has been observed for other ALS/FTD-associated RBPs, in which the ability to bind RNAs is a key mediator of overexpression toxicity.

### The toxicity of RNA binding-deficient MATR3 variants is highly dependent on their subcellular distribution

One of the hallmarks of neurodegenerative diseases, including ALS and FTD, is the formation of protein-rich aggregates (Arai et al., 2006; Neumann et al., 2006). Prior investigations suggest that these aggregates may be toxic, innocuous, or representative of a coping response that ultimately prolongs neuronal survival (Arrasate et al., 2004; Barmada et al., 2010). To determine if the formation of nuclear puncta by MATR3(dRRM2)-EGFP and MATR3(dRRM1/2)-EGFP affected neuronal lifespan, we turned to LFM. We employed a modified version of the automated analysis script to draw ROIs around the nuclear perimeter within each transfected cell (Fig. 4A) and then calculated a coefficient of variation (CV) for the MATR3(dRRM1/2)-EGFP signal within each nuclear ROI. The CV, or the ratio of the standard deviation of GFP intensity to the mean GFP intensity for the ROI, is directly proportional to the spatial variability of fluorescence intensity within each ROI. Therefore, we reasoned that this measure might be useful for rapidly and reliably identifying puncta in an unbiased and high-throughput manner. We first validated the use of CV for detecting puncta by creating a receiver-operator characteristic (ROC) curve; in doing so, we observed that a CV threshold of 0.92 was 87.2% sensitive and 93.9% specific in discriminating cells with nuclear granules from those with diffuse protein (Fig. 4B). We therefore utilized this CV threshold to assess the frequency of nuclear granule formation in primary rodent cortical neurons, noting that 24 h after transfection, 23.4% (653/2734) of neurons transfected with MATR3(dRRM2)-EGFP neurons displayed nuclear granules compared to only 8.8% (153/1743) of MATR3(dRRM1/2)-EGFP cells (Fig. 4C). We also observed the time-dependent formation of nuclear granules as neurons expressed increasing amounts of MATR3-EGFP (Fig. 4D), suggesting that granule formation may be proportional to expression level. To investigate this relationship further, we identified neurons exhibiting a diffuse distribution of MATR3(dRRM2)-EGFP at day 1 and followed these cells for an additional 3 days by automated microscopy. We then measured the GFP intensity for each cell at day 1, and related this value to the risk of granule formation over the ensuring 72 h period using penalized splines models. Notably, we failed to observe a significant relationship between GFP intensity on day 1 and granule formation by day 3 (Fig. 3E). We also assessed the relative change in expression level on a per-cell basis, as quantified by the ratio of GFP intensity at day 2 to the GFP intensity at day 1, to determine if the net rate of MATR3(dRRM2)-EGFP production better predicted granule formation. The probability of granule formation was directly proportional to the time-dependent change in MATR3(dRRM2)-EGFP levels (Fig. 4F), suggesting that granule formation is favored by the rapid accumulation of MATR3(dRRM2)-EGFP.

**Figure 4.**
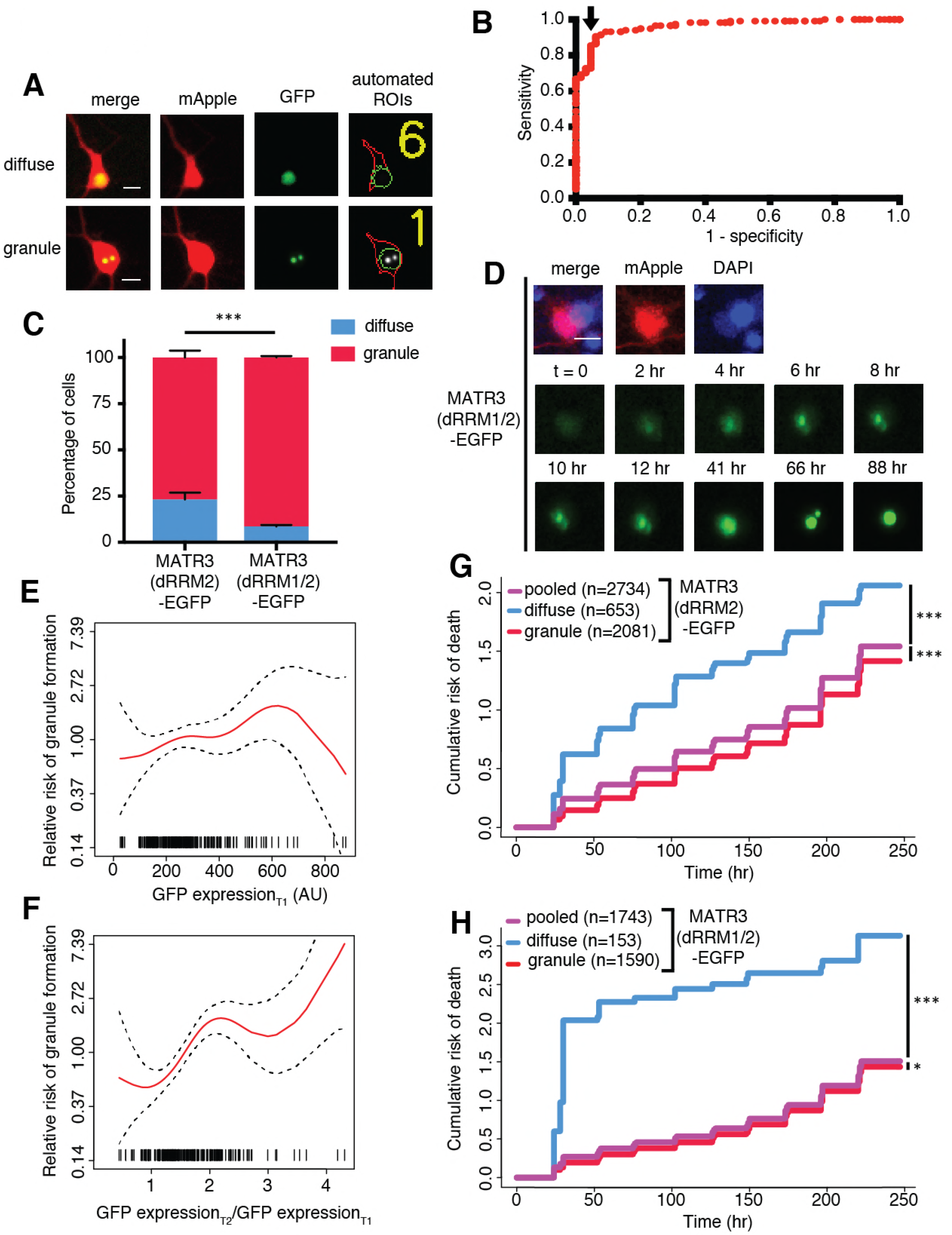
MATR3(dRRM2)-EGFP and MATR3(dRRM1/2)-EGFP are highly neurotoxic in their diffuse form. **A.** Automated analysis of MATR3-EGFP distribution in transfected primary cortical neurons. Regions of interest (ROIs) were drawn around the cell body (marked by mApple fluorescence, red) and diffuse MATR3-EGFP (indicated by GFP fluorescence, green), and used to calculate a coefficient of variation (CV) representing MATR3-EGFP distribution within each ROI. **B.** Receiver operating characteristic (ROC) curve for MATR3-EGFP CV values. A CV threshold of 0.92 (arrow) identified cells with intranuclear MATR3-EGFP granules with 87.2% sensitivity and 93.9% specificity. **C.** Using this cutoff, we determined that 1 day after transfection, 23.4% (653/2734) of MATR3(dRRM2)-EGFP neurons displayed intranuclear granules compared to only 8.8% (153/1743) of MATR3(dRRM1/2)-EGFP cells. (p < 0.00001; Fisher’s exact test). **D.** Intranuclear granules form in a time-dependent manner in neurons expressing MATR3(dRRM2)-EGFP and MATR3(dRRM1/2)-EGFP. **E-F.** Penalized spline models depicting the relationship between MATR3(dRRM2)-EGFP expression on day 1 (**E**) or change in GFP expression between day 1 and day 2 (**F**), and risk of developing an intranuclear granule by day 3. Expression level at day 1 was not significantly associated with risk of granule formation (**E**; p = 0.30; penalized spline regression), but the relative increase in expression from day 1 to day 2 is (**F**; p = 0.015; penalized spline regression). **G.** For MATR3(dRRM2)-EGFP, neurons exhibiting granules by day 1 displayed improved survival compared to the pooled combination of all cells. Conversely, neurons with diffusely distributed MATR3(dRRM2)-EGFP fared far worse (comparing to the pooled condition: cells with granules n = 2081, HR = 0.86, p = 1.02 × 10^−5^; cells with diffuse protein n = 653, HR = 1.75, p < 2 × 10“^16^; Cox proportional hazards). **H.** Neurons with MATR3(dRRM1/2)-EGFP granules by day 1 similarly displayed a reduced risk of death in comparison to the pooled group, while diffuse MATR3(dRRM1/2)-EGFP was highly toxic (comparing to the pooled condition: cells with granules n = 1590, HR = 0.92, p = 0.03; cells with diffuse protein n = 153, HR = 3.78, p = 2 × 10^−16^; Cox proportional hazards). Scale bars in (**A**) and (**B**), 10 μm.

Our previous studies demonstrated that deletion of RRM1 or RRM1 and 2 had no effect upon the toxicity of MATR3-EGFP when expressed in primary neurons (Fig. 3E). These analyses included all neurons within a given condition, consisting of cells with diffuse nuclear MATR3 as well as those with MATR3 redistributed into granules. To determine if the presence of nuclear MATR3-EGFP granules impacted the survival of neurons, we utilized the nuclear CV threshold (Fig. 4B) to divide neurons expressing MATR3(dRRM2)-EGFP and MATR3(dRRM1/2)-EGFP into three categories: cells with diffuse protein at day 1, those with granules at day 1, or all cells. We then tracked neurons in each category for the following 9 days by LFM, and compared their survival by Cox proportional hazards analysis. By these measures, neurons displaying nuclear MATR3(dRRM2)-EGFP granules fared significantly better than the population as a whole, while those exhibiting a diffuse distribution demonstrated an increased risk of death (Fig. 4G). Similar results were obtained for neurons expressing MATR3(dRRM1/2)-EGFP; here, the relative protection associated with nuclear MATR3(dRRM1/2)-EGFP granules was modest, but the toxicity of diffusely-distributed MATR3(dRRM1/2)-EGFP was more pronounced (Fig. 4H). The marked toxicity of diffuse MATR3(dRRM1/2)-EGFP may explain why so few cells with diffuse protein are seen at day 1 (Fig. 4D). Taken together, these results suggest that diffuse MATR3 is highly neurotoxic when it cannot bind RNA. Furthermore, the sequestration of RNA binding-deficit MATR3 variants into nuclear granules is associated with a survival advantage.

### MATR3 granules formed by deletion of the RNA-binding domains display liquidlike properties that are affected by pathogenic mutations

As part of their normal function, many RBPs reversibly undergo liquid-liquid phase separation (LLPS), involving the formation of droplets with liquid-like properties from diffuse or soluble proteins (Molliex et al., 2015; Murray et al., 2017). Disease-associated mutations in the genes encoding these proteins may promote LLPS or impair the reversibility of phase separation (Molliex et al., 2015; Patel et al., 2015; Conicella et al., 2016). We wondered whether the intranuclear granules formed by MATR3(dRRM2)-EGFP and MATR3(dRRM1/2)-EGFP represent liquid droplets and also whether pathogenic MATR3 mutations affect the intrinsic properties of these puncta. Indeed, nuclear granules exhibited dynamic properties, not only growing in size over time but also moving freely within the nucleus and fusing if they encountered other granules (Fig. 5A), indicative of liquid-like behavior.

**Figure 5.**
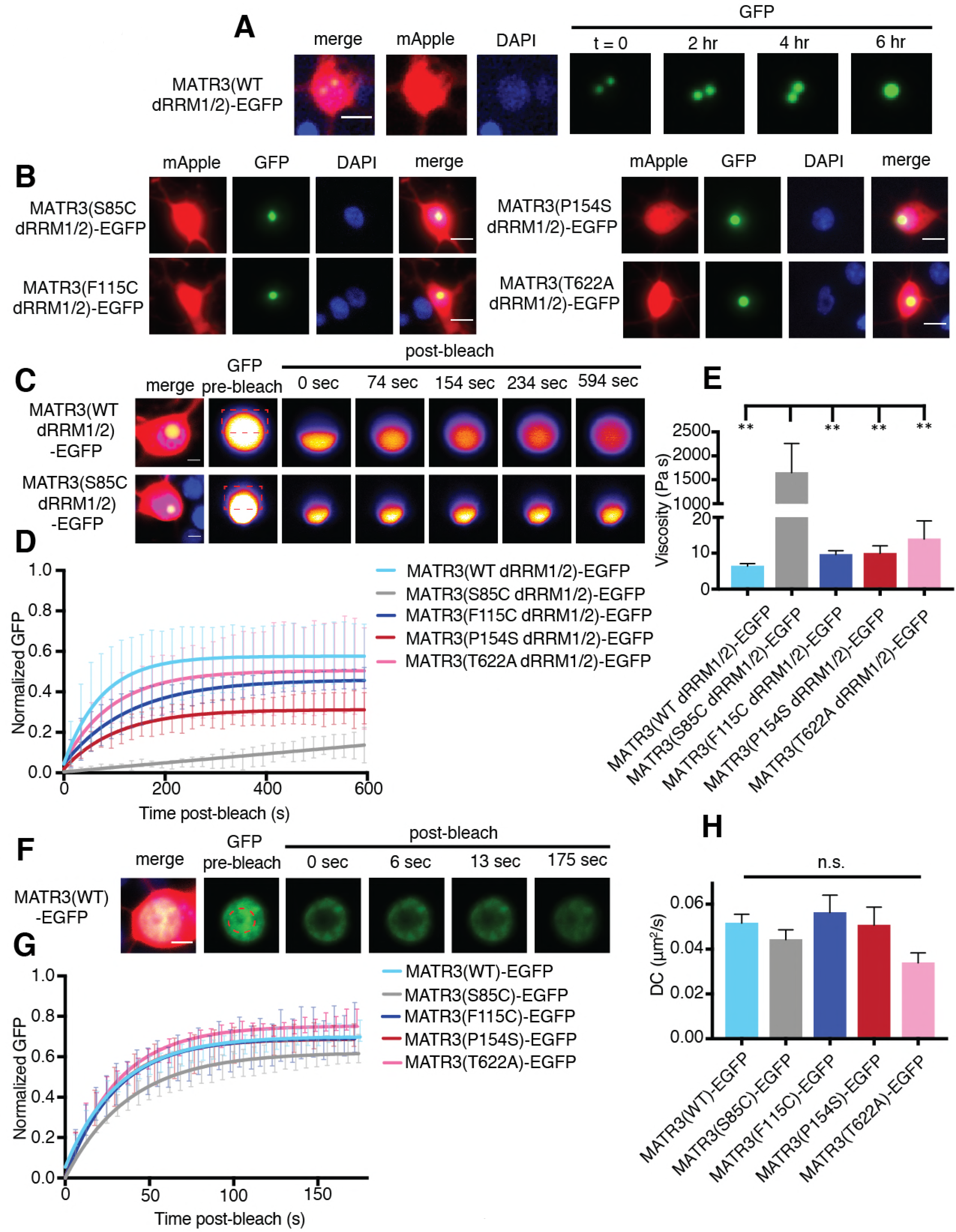
MATR3(dRRM1/2)-EGFP droplets display liquid-like properties that are affected by the S85C mutation. **A.** MATR3(dRRM1/2)-EGFP and MATR3(dRRM1/2)-EGFP droplets show liquid-like properties such as mobility and fusion. **B.** Pathogenic MATR3 mutations on the dRRM1/2 background result in similar phase-separated droplets. **C-D.** Fluorescence recovery after photobleaching (FRAP) of MATR3(dRRM1/2)-EGFP droplets shows internal rearrangement consistent with liquidlike behavior, but the recovery of MATR3(S85C dRRM1/2)-EGFP droplets was significantly delayed. **E.** MATR3(S85C dRRM1/2)-EGFP droplets displayed significantly higher viscosity in comparison to other variants (comparing to S85C MATR3(dRRM1/2)-EGFP n = 5: WT MATR3(dRRM1/2)-EGFP n = 5, p = 0.0045; F115C MATR3(dRRM1/2)-EGFP n = 5, p = 0.0046; P154S MATR3(dRRM1/2)-EGFP n = 5, p = 0.0046; T622A MATR3(dRRM1/2)-EGFP n = 4, p = 0.0079; one-way ANOVA with Tukey’s post-hoc test). **F-G.** FRAP experiments involving full-length MATR3-EGFP variants showed no differences in rates of return. **H.** Similarly, there were no differences in diffusion coefficients (DC) among full-length MATR3 variants (MATR3(WT)-EGFP n = 5, MATR3(S85C)-EGFP n = 5, MATR3(F115C)-EGFP n = 5, MATR3(P154S)-EGFP n = 5, MATR3(T622A)-EGFP n = 4); p = 0.17; one-way ANOVA). Scale bars in (**A**) and (**B**), 10 μm; scale bars in (**C**) and (**F**), 5 μm.

We then asked if these structures displayed internal rearrangement characteristic of liquid droplets (Lin et al., 2015; Shin and Brangwynne, 2017) and whether pathogenic *MATR3* mutations affect their dynamics. To answer this, we introduced disease-associated mutations into MATR3(dRRM1/2)-EGFP, and transfected rodent primary cortical neurons with each construct (Fig. 5B). Nuclear puncta were photobleached 2-4 days after transfection, and the recovery of fluorescence intensity tracked within the bleached and unbleached ROIs by laser scanning confocal microscopy. Granules formed by WT MATR3(dRRM1/2)-EGFP displayed internal rearrangement over the course of minutes consistent with liquid-like properties, as did all tested disease mutants on the dRRM1/2 background (Fig. 5C-D). The S85C mutation, however, severely slowed fluorescence recovery, suggesting reduced exchange of molecules within each droplet. Using the Stokes-Einstein equation, we calculated viscosity estimates for each MATR3(dRRM1/2)-EGFP variant based on return time and bleached area size (Fig. 5E). Consistent with the observed effect of this mutation on fluorescence recovery, the S85C mutation led to a pronounced increase in viscosity over that of WT and other disease-associated mutants.

We wondered whether this phenotype was specific to nuclear droplets formed by MATR3(dRRM1/2)-EGFP, or if full-length MATR3 carrying pathogenic mutations would also display reduced mobility. For this, we transfected primary neurons with full-length versions of MATR3(WT)-EGFP or disease-associated MATR3-EGFP variants and then bleached a circular area in the center of the nucleus (Fig. 5F). In each case, we noted rapid return of fluorescence, and the recovery rate was unaffected by pathogenic *MATR3* point mutations (Fig. 5G). To account for the rapidity of return as well as the area of the bleached region, we calculated a diffusion coefficient (DC) for each construct. Comparison of the DCs for WT and mutant MATR3-EGFP variants showed no significant differences (Fig. 5H). Our data therefore suggest that the S85C point mutation—and perhaps other mutations that cluster in the N-terminal disordered domain—selectively affect the droplet properties of MATR3.

### Mapping the sequence determinants of MATR3 localization in neurons

Cytoplasmic inclusions composed of the RBP TDP-43 are characteristic of ALS and the majority of FTD (Arai et al., 2006; Neumann et al., 2006). Moreover, pathogenic mutations in the gene encoding TDP-43 enhance cytoplasmic mislocalization concordant with enhanced neurotoxicity, and reductions in cytoplasmic TDP-43 prolong neuronal survival (Barmada et al., 2010; Barmada et al., 2014). To determine if MATR3 localization is likewise an important determinant of neurodegeneration, we sought to disrupt the MATR3 nuclear localization signal (NLS). However, since multiple sequences have been associated with nuclear MATR3 localization (Hibino et al., 2006; Hisada-Ishii et al., 2007), we systematically identified regions enriched in positively-charged amino acids (arginine, lysine) that may mediate nuclear import via importin-α. We then deleted each of the 7 regions defined in this manner, including two that had been identified as controlling nuclear localization in previous studies, and assessed their localization by transfection in rodent primary cortical neurons followed by fluorescence microscopy (Fig. 6A).

**Figure 6.**
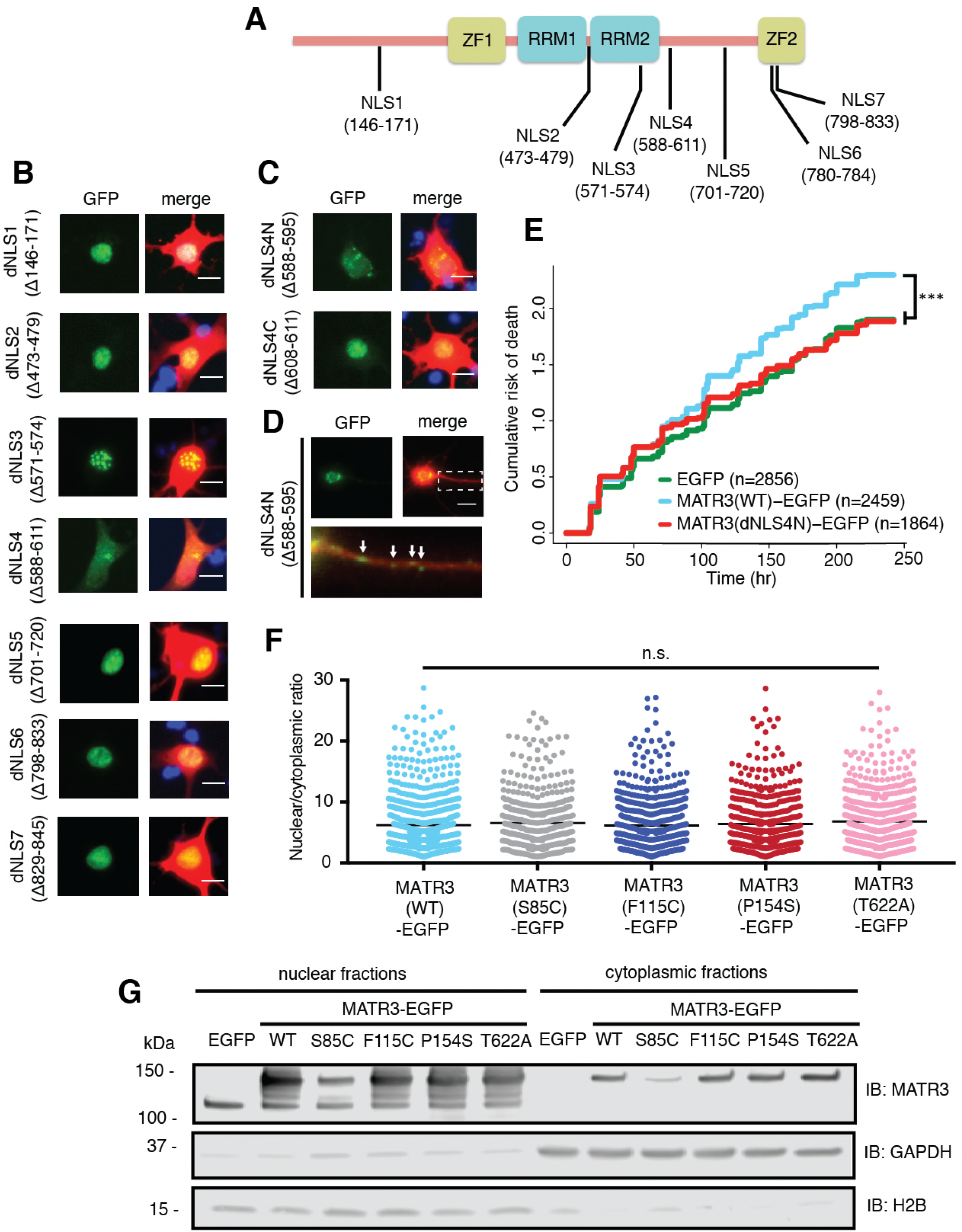
Reducing MATR3 nuclear localization mitigates overexpression toxicity. **A.** Schematic showing putative MATR3 nuclear localization signals (NLS). **B-C.** Deletion of the N-terminal arm of NLS4 (dNLS4N) led to nuclear MATR3 clearance in neurons. **D.** MATR3(dNLS4N)-EGFP forms granular structures in the cytoplasm and neuronal processes (white arrows). **E.** Disrupting nuclear localization of MATR3 prevents neurotoxicity from overexpression (MATR3(WT)-EGFP n = 2459; MATR3(dNLS4N)-EGFP n = 1864, HR = 0.89, p = 0.00041; Cox proportional hazards). **F-G.** Pathogenic MATR3 mutants display no difference in subcellular protein localization as assessed by automated image nuclear/cytoplasmic analysis (**F**; MATR3(WT)-EGFP n = 824, MATR3(S85C)-EGFP n = 499, MATR3(F115C)-EGFP n = 634, MATR3(P154S)-EGFP n = 554, MATR3(T622A)-EGFP n = 677; p = 0.067; one-way ANOVA) or biochemical fractionation in transfected HEK293T cells (**G**). Nevertheless, Western blot demonstrated reduced abundance of the S85C mutant in transfected HEK293T cells. Scale bars in (**B**) and (**C**), 10 μm; scale bar in (D), 50 μm.

Deletion of NLSs 1, 2, 3, 5, 6, and 7 had little to no effect on neuronal MATR3 distribution (Fig. 6B). While the dNLS3 mutation did not change nuclear MATR3 localization *per se*, it did induce the formation of many small, nuclear granules. This effect is consistent with the position of NLS3 within RRM2, and the observed formation of nuclear puncta upon RRM2 deletion (Fig. 4). In contrast, and in accord with previous studies (Hisada-Ishii et al., 2007), deletion of the bipartite NLS4 elicited a marked reduction in nuclear MATR3-EGFP accompanied by enhanced cytoplasmic localization and the formation of small MATR3-EGFP granules within the cytoplasm. In DT40 and HeLa cells, both NLS4 arms were critical for MATR3 nuclear localization (Hisada-Ishii et al., 2007). To determine if this is the case in neurons, we sequentially deleted the N- and C-terminal arms (dNLS4N and dNLS4C, respectively) and tested their localization by transfection into primary cortical neurons. These studies demonstrated that the N-terminal arm is necessary and sufficient for nuclear localization, as MATR3(dNLS4N)-EGFP exhibits nuclear clearing and punctate distribution in the cytoplasm and neuronal processes, while MATR3(dNLS4C)-EGFP has the same distribution as MATR3(WT)-EGFP (Fig. 6C-D).

Having identified the N-terminal arm of NLS4 as the key sequence regulating MATR3 localization in neurons, we asked whether driving MATR3 into the cytoplasm by deletion of this sequence could modify toxicity. Rodent primary cortical neurons were transfected with mApple and either EGFP, MATR3(WT)-EGFP, or MATR3(dNLS4N)-EGFP and imaged at regular intervals by LFM. Automated survival analysis of neuronal populations expressing these constructs demonstrated that the dNLS4N mutation and resulting cytoplasmic localization significantly reduced MATR3-dependent toxicity compared to the MATR3(WT)-EGFP (Fig. 6E). Therefore, unlike TDP-43 and FUS, two RBPs whose cytoplasmic mislocalization are tightly tied to neurodegeneration in ALS/FTD models, cytoplasmic MATR3 retention mitigates toxicity, suggesting that nuclear MATR3 functions are required for neurodegeneration (Barmada et al., 2010; Qiu et al., 2014).

Given the observed relationship between MATR3 localization and toxicity, we wondered if subtle changes in nucleocytoplasmic MATR3 distribution could be responsible for the increased toxicity of MATR3 bearing disease-associated mutations. Rodent primary cortical neurons transfected with each of the pathogenic MATR3-EGFP variants showed no obvious difference in subcellular localization in comparison with MATR3(WT)-EGFP (Fig. 2A). To investigate MATR3-EGFP localization in a quantitative manner, we developed a customized image-based analysis script to draw ROIs around the nucleus and soma of each neuron, measure MATR3-EGFP content separately within each compartment, and calculate a nucleocytoplasmic ratio for MATR3-EGFP in individual cells (Fig. 6F). This analysis confirmed our initial observations, showing no significant differences in the localization of mutant MATR3-EGFP variants compared to MATR3(WT)-EGFP.

In a complementary series of experiments, we utilized biochemical fractionation to assess the distribution of MATR3-EGFP in a human cell line. MATR3(WT)-EGFP or versions of MATR3-EGFP bearing the S85C, F115C, P154S, and T622A disease-associated mutations were transfected into HEK293T cells, and the nuclear and cytoplasmic fractions subjected to SDS-PAGE and Western blotting. In agreement with single-cell data from transfected primary neurons, we noted no difference in the nucleocytoplasmic distribution of any of the MATR3-EGFP variants tested here (Fig. 6G). Nevertheless, we consistently observed far less of the S85C variant in both nuclear and cytoplasmic fractions, compared to MATR3(WT)-EGFP and other disease-associated mutants. These data suggest that the S85C mutation may destabilize MATR3-EGFP; alternatively, this mutation may prevent adequate solubilization and detection of MATR3-EGFP via SDS-PAGE and Western blotting.

### A subset of pathogenic MATR3 mutations affect protein solubility but not stability

To discriminate among these possibilities, we first investigated the turnover of WT and mutant MATR3 variants using optical pulse labeling (OPL), a technique enabling non-invasive determinations of protein clearance in living cells (Barmada et al., 2014). For these experiments, MATR3 was fused to Dendra2—a photoconvertable protein that irreversibly switches from a green to red fluorescent state upon illumination with low-wavelength light (Chudakov et al., 2007)—and expressed in primary cortical neurons. One day after transfection, neurons were illuminated with blue light to photoconvert Dendra2, and the time-dependent loss of red fluorescence signal used to calculate protein half-life (Fig. 7A). Previous studies validated the accuracy and utility of OPL for determinations of protein half-life (Barmada et al., 2014); importantly, and in contrast to biochemical techniques for calculating half-life that depend on radioactive labeling or translational inhibitors, OPL allows us to measure protein clearance on a single-cell level for thousands of neurons simultaneously (Fig. 7B). Most disease-associated mutations had little effect upon the turnover of MATR3-Dendra2 in primary cortical neurons. However, we noted subtle destabilization of MATR3(S85C)-Dendra2 in comparison to other pathogenic mutant variants and MATR3(WT)-Dendra2 (Fig. 7C-D). Even so, the magnitude of the effect was relatively small, making it unlikely that differences in protein turnover fully explain the reduced abundance of MATR3(S85C)-EGFP noted in cell lysates (Fig. 6G).

**Figure 7.**
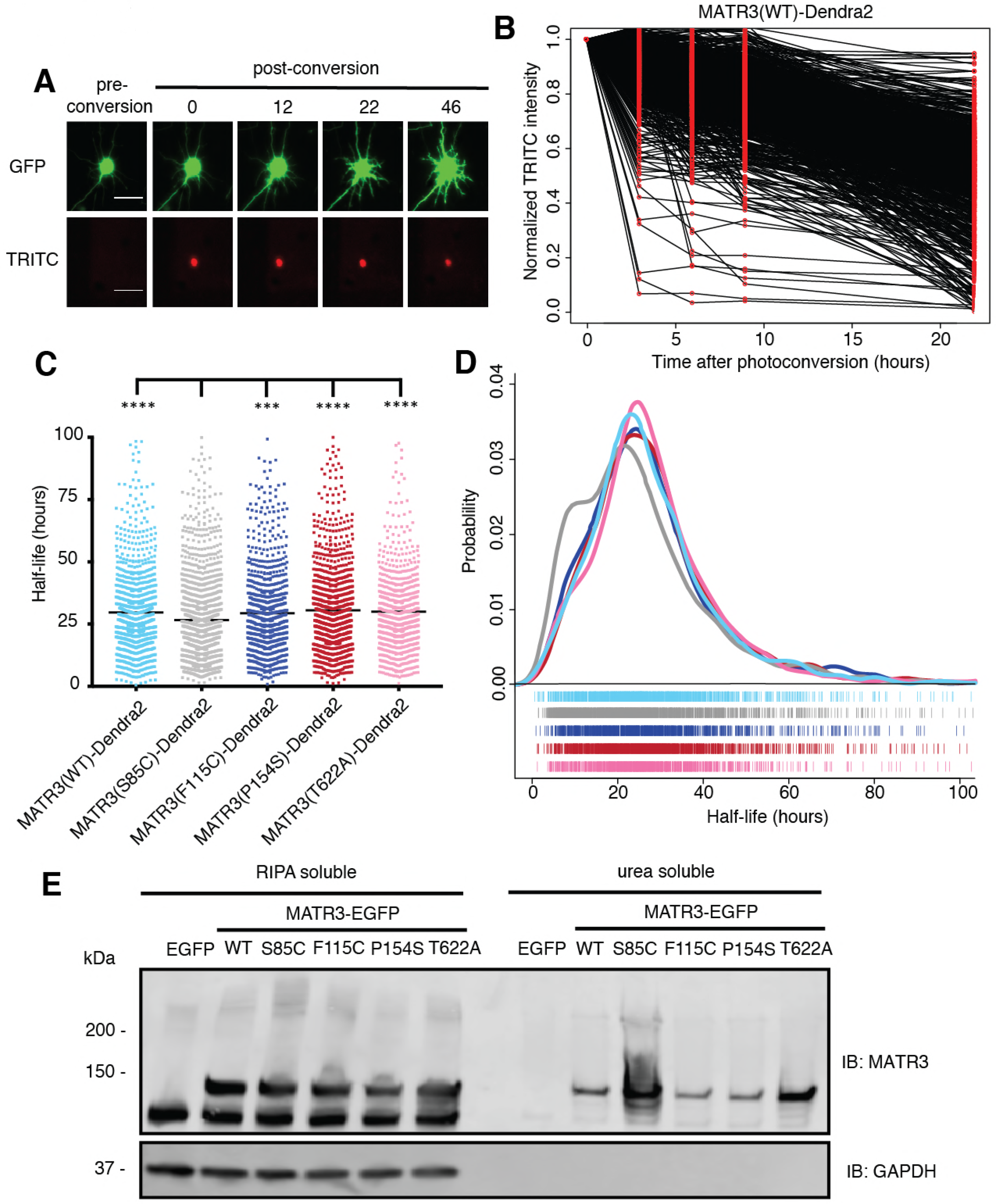
Pathogenic MATR3 mutations have little effect on MATR3 turnover, but a subset reduce solubility. **A.** Optical pulse labeling of Dendra2-tagged MATR3 variants. Each neuron is transfected with EGFP alone to outline the cell body, as well as MATR3-Dendra2, which fluoresces in the red channel (TRITC) upon photoconversion. Scale bar, 50 μm. **B.** Normalized red fluorescence (TRITC) signal for individual neurons. The time-dependent decay of red fluorescence over time is used to calculate MATR3-Dendra2 half-life for each neuron. **C-D.** MATR3(S85C)-Dendra2 displayed a subtle but significant reduction in half-life compared to MATR3(WT)-Dendra2 and the other pathogenic mutants tested (comparing to MATR3(S85C)-Dendra2 n = 1670: MATR3(WT)-Dendra2, n = 1269, p < 0.0001; MATR3(F115C)-Dendra2, n = 1122, p = 0.0001; MATR3(P154S)-Dendra2, n = 1509, p < 0.0001; MATR3(T622A)-Dendra2, n = 923, p < 0.0001; one-way ANOVA with Tukey’s post-hoc test). **E.** Sonication in RIPA resulted in equivalent amounts of all MATR3 variants by Western blotting. The S85C variant was markedly enriched in the RIPA-insoluble, urea-soluble fraction, while the T622A variant showed more modest enrichment.

We next asked if the S85C mutation altered MATR3 solubility. HEK293T cells transfected with WT and mutant MATR3-EGFP variants were lysed using a harsher protocol that involved sonication in RIPA buffer; additionally, we used urea buffer to extract all RIPA-insoluble proteins. In stark contrast to mild conditions (Fig. 6G), harsher lysis resulted in equivalent levels of all MATR3 variants on Western blot, suggesting that the S85C mutation reduced MATR3 solubility (Fig. 7E). Consistent with this interpretation, the urea-soluble fraction was markedly enriched for MATR3(S85C)-EGFP and modestly enriched for MATR3(T622A)-EGFP. These data show that the S85C and T622A mutations reduce the solubility of MATR3, without drastically affecting its stability. As shown in Fig. 1A, both mutations lie within areas of predicted disorder, consistent with their effects on MATR3 aggregation and solubility.

## DISCUSSION

In this study, we modeled MATR3-mediated neurodegeneration by overexpressing WT or disease-associated MATR3 variants in primary neurons. In doing so, we found that neurons were highly susceptible to increases or decreases in MATR3 levels, and disease-associated MATR3 variants exhibited enhanced toxicity in comparison to MATR3(WT). Structure-function studies demonstrated that the ZF2 domain modulates overexpression-related toxicity, while RRM2 prevents MATR3 phase separation into mobile nuclear puncta. Biophysical analysis of these puncta confirmed their liquid-like nature and further indicated that the pathogenic S85C mutation substantially increased the viscosity of these structures. We also determined that the N-terminal arm of a bipartite NLS drives MATR3 nuclear localization; forcing MATR3 into the cytoplasm by deleting this sequence blocked toxicity from MATR3 overexpression. While we did not observe any differences in the distribution of pathogenic MATR3 variants, we noted that the S85C mutation significantly reduced MATR3 solubility and, to a lesser extent, stability. The T622A mutant displayed similar but more muted effects on MATR3 solubility, suggesting that disease-associated mutations located in distinct MATR3 domains may operate through convergent pathogenic mechanisms.

Both MATR3 overexpression and knockdown elicited significant and comparable toxicity in neurons. These data suggest that neurons are bidirectionally vulnerable to changes in MATR3 levels. Post-mortem studies of MATR3 distribution in sporadic and familial ALS patients demonstrated stronger MATR3 nuclear staining as well as the presence of cytoplasmic MATR3 aggregates in motor neurons (Dreser et al., 2017; Tada et al., 2017). While the impact of these findings is unknown, MATR3 mislocalization or sequestration into aggregates may reflect a reduction in normal function, a new and abnormal function, or both. In mice, homozygous *Matr3* knockout is embryonic lethal, while heterozygous *Matr3*^+/-^ animals demonstrate incompletely penetrant cardiac developmental abnormalities. However, *Matr3*^+/−^ mice exhibited roughly equivalent Matr3 protein levels in comparison to nontransgenic animals, complicating any conclusions regarding Matr3 loss-of-function in these models (Quintero-Rivera et al., 2015). Overexpression of human MATR3(F115C) in mice results in severe muscle disease consisting of fore-and hindlimb muscle atrophy accompanied by vacuolization (Moloney et al., 2016). These animals also displayed spinal cord gliosis and cytoplasmic MATR3 redistribution in spinal motor neurons akin to changes in MATR3 localization noted in humans with ALS, although no significant neurodegeneration was observed in MATR3(F 115C) transgenic mice. Our data illustrating the dose-dependency of MATR3 neurotoxicity (Fig. 1) imply that MATR3(F115C) expression may be insufficient to elicit neurodegeneration in these animals. Alternatively, constitutive overexpression of MATR3(F115C) in transgenic mice may trigger compensatory mechanisms during development that promote neuronal survival.

MATR3 is unique among ALS/FTD-associated RBPs in possessing not just two tandem RRMs but also two ZF domains that can bind repetitive DNA elements found in the nuclear scaffold, consistent with MATR3’s localization within the nuclear matrix (Hibino et al., 1998). We attempted to identify which functional domains were important for MATR3 overexpression toxicity and found that while deletion of ZF2 resulted in modest rescue, deletion of RRM2—either alone or in combination with RRM1—resulted in the formation of phase-separated intranuclear droplets. Our data therefore support a model in which RNA binding prevents MATR3 self-association into droplets. Consistent with this interpretation, we observed small, mobile MATR3 granules in the cytoplasm and neuronal processes when the bipartite NLS was disrupted (Fig. 6D). Cytoplasmic RNA concentrations are more than an order of magnitude lower than those in the nucleus, a gradient that may favor the coalescence of MATR3(dNLS4N)-EGFP into puncta within the neuronal soma and processes (Goldstein and Trescott, 1970).

The functional importance of the individual RRM domains for MATR3’s RNA binding activity is unclear; while some studies suggest that both RRM1 and RRM2 bind RNA, other investigations indicated that RRM2 is primarily responsible for binding RNA (Hibino et al., 2006; Salton et al., 2011). Our data show that deletion of RRM2 is sufficient to elicit phase separation by MATR3, suggesting that RNA recognition by MATR3 is mediated largely by RRM2. We also noted no significant difference in the survival of neuronal populations overexpressing dRRM1, dRRM2, and dRRM1/2 variants of MATR3-EGFP, implying that RNA binding *per se* is unrelated to MATR3-mediated neurodegeneration. This interpretation is strengthened by detailed analyses of neurons expressing MATR3(dRRM2) and MATR3(dRRM1/2). When neurons with and without droplets were assessed separately, we noted that neurons exhibiting diffuse MATR3(dRRM2) or MATR3(dRRM1/2) displayed a significantly higher risk of death than those with droplets. These results imply that diffuse MATR3, when not bound to RNA, can be highly toxic. Conversely, sequestration of RNA-binding deficient MATR3 into puncta is associated with extended neuronal survival. Our data further indicate that diffuse MATR3(dRRM1/2) is more toxic than diffuse MATR3(dRRM2) (compare the diffuse population in Fig. 4G to the diffuse population in Fig. 4H). Since RRM1 may be capable of recognizing some RNA even without RRM2, these observations suggest that neurodegeneration is inversely proportional to the ability of MATR3 to bind RNA when diffusely localized within the nucleus. In disease models involving related RBPs, including TDP-43 and FUS, toxicity requires the presence of RNA binding motifs as well as low-complexity domains that enable LLPS (Johnson et al., 2008; Daigle et al., 2013; Ihara et al., 2013). As with MATR3, abrogation of RNA binding may disinhibit self-association, resulting in the sequestration of otherwise toxic diffuse protein within droplets.

Investigating the liquid-like properties of MATR3(dRRM1/2)-EGFP droplets, we noted a selective effect of the S85C mutation on droplet viscosity. Low-complexity, intrinsically disordered domains are required for phase separation and self-assembly of RBPs. Apart from its nucleic acid binding domains, MATR3 displays a high degree of predicted disorder based on its primary amino acid sequence (Fig. 1A). The location of the S85C mutation and its effects on MATR3(dRRM2)-EGFP droplet viscosity suggest that the N-terminal disordered region of MATR3 regulates the liquid-like properties of droplets. Whether full-length MATR3 is capable of phase-separation under physiological circumstances, and what relevance this process has for disease, is currently unclear.

Conflicting evidence (Hibino et al., 2006; Hisada-Ishii et al., 2007) suggests that MATR3 nuclear import is driven by distinct sequences in different cell types. For example, while amino acids 701-718 are essential for nuclear localization of rat MATR3 in Ac2F cells, deletion of the homologous sequence (amino acids 701-720) in human MATR3 has no effect on neuronal distribution (Fig. 6B). To identify the sequences responsible for MATR3 nuclear import within neurons, we undertook a systematic analysis of arginine/lysine-rich sequences in MATR3 resembling NLSs. In accord with an earlier report (Hisada-Ishii et al., 2017), we found that MATR3’s bipartite NLS (NLS4) controlled its nuclear enrichment in neurons, but only the N-terminal arm of the NLS was sufficient for MATR3 nuclear clearing and cytoplasmic distribution. Pathogenic *TARDBP* and *FUS* mutations promote cytoplasmic mislocalization of TDP-43 and FUS, respectively, and cytoplasmic enrichment of these proteins is tightly linked to toxicity (Barmada et al., 2010; Dormann et al., 2010). In stark contrast, however, we observed that cytoplasmic MATR3 redistribution extended neuronal survival, suggesting—along with the partial rescue we observed for MATR3(dZF2)-EGFP and MATR3(dZF1/2)-EGFP—that MATR3 overexpression elicits neurodegeneration through nuclear DNA binding activity, mediated at least in part by ZF2.

Given previously established relationships between the distribution and aggregation of RBPs and neurodegeneration in ALS models (Johnson et al., 2009; Barmada et al., 2010; Dormann et al., 2010; Igaz et al., 2011; Kim et al., 2013; Qiu et al., 2014), we wondered whether the enhanced toxicity of pathogenic MATR3 variants arises from mutation-associated changes in MATR3 localization or solubility. We noted no significant differences in the subcellular distribution of mutant MATR3 variants in comparison to MATR3(WT), but instead consistently observed reduced levels of MATR3(S85C) in transfected cell lysates. A similar pattern was noted in previous investigations and attributed to reduced MATR3(S85C) stability (Johnson et al., 2014). Using OPL, a sensitive method for measuring protein turnover *in situ* (Barmada et al., 2014; Gupta et al., 2017), we detected only a very modest shortening of MATR3(S85C) half-life compared to MATR3(WT). Nevertheless, we observed a marked change in the solubility of MATR3(S85C) and, less so, MATR3(T622A). This is in partial agreement with initial studies of MATR3(S85C) that noted equivalent amounts of MATR3(WT) and MATR3(S85C) in insoluble fractions but reduced MATR3(S85C) in the nuclear fraction (Senderek et al., 2009). Both the S85C and T622A mutations lie within domains predicted to be disordered (Fig. 1). Furthermore, both mutations disrupt potential phosphorylation sites, and phosphorylation within the intrinsically disordered domain of FUS inhibits self-association of the protein through negative-negative charge repulsion between phosphate groups (Monahan et al., 2017). Of the 13 pathogenic mutations identified to date in MATR3, four (S85C, S610F, T622A, S707L) eliminate phosphorylatable residues, suggesting that inadequate phosphorylation and subsequent disinhibited self-association of MATR3 may be a conserved feature of MATR3 mutants.

MATR3’s possesses broad functions in DNA/RNA processing (Belgrader et al., 1991; Hibino et al., 2000; Zhang and Carmichael, 2001; Salton et al., 2014; Coelho et al., 2015; Rajgor et al., 2016; Uemura et al., 2017). Its presence within cytoplasmic aggregates in approximately half of patients with sporadic ALS (Tada et al., 2017) implies that MATR3 pathology causes or is caused by cellular alterations in RNA and protein homeostasis, many of which may contribute to neurodegeneration in ALS and related disorders. Our work confirms that MATR3 is essential for maintaining neuronal survival and furthermore shows that MATR3 accumulation results in neurodegeneration in a manner that depends on its subcellular localization and ZF domains. Additional studies are required to further delineate the impact of disease-associated MATR3 mutations on the function, behavior, and liquid-like properties of MATR3.

## MATERIALS AND METHODS

### Plasmids

Full-length human *MATR3* cDNA was obtained from Addgene (#32880) and cloned into the pCMV-Tag2B vector (Agilent Technologies, #211172, Santa Clara, CA) using BamHI and XhoI endonucleases, tagging the amino-terminus with a FLAG epitope. To generate MATR3-EGFP, the *EGFP* open reading frame with a 14 amino acid N-terminal linker was amplified from pGW1-EGFP (Arrasate et al., 2004) by PCR using forward primer AGC TAC TAG TAC TAG AGC TGT TTG GGA C and reverse primer TAT TGG GCC CCT ATT ACT TGT ACA GCT CGT CCA T. The resulting amplicon was digested with SpeI and ApaI and cloned into the corresponding sites in pKS to generate pKS-EGFP. To create pKS-MATR3-EGFP, the *FLAG-MATR3* open reading frame from pCMV-Tag2B was amplified by PCR with forward primer GAT CTC TAG AGC GGC CGC CAC CAT GGA T and reverse primer AGC TAC TAG TCA TAG TTT CCT TCT TCT GTC T, digested with XbaI and SpeI, and inserted into the corresponding sites in pKS-EGFP. pGW1-MATR3-EGFP was generated by digesting pKS-MATR3-EGFP with XbaI and ApaI, purifying the ensuing fragment containing MATR3-EGFP, and inserting into the corresponding sites of pGW1. To create Dendra2-tagged MATR3 variants, the *EGFP* coding region of each construct was removed by PCR amplification of the pGW1-MATR3-EGFP vector using primers that flank the *EGFP* open reading frame. The *Dendra2* open reading frame was then removed from pGW1-Dendra2 (Barmada et al., 2014) by digestion with ApaI and MfeI, and inserted into pGW1-MATR3. All constructs were confirmed by sequencing prior to transfection in neurons and HEK293T cells.

Domain deletion mutants were created using Q5 Hot Start High-Fidelity DNA Polymerase (New England Biolabs, Ipswich, MA) and primers flanking the regions to be deleted for nucleic acid-binding domain (Table 1) and putative nuclear localization signal (Table 2) deletions. All disease-associated point mutations were created with site-directed mutagenesis (Table 3).

### Primary neuron cell culture and transfection

Cortices from embryonic day (E)19-20 Long-Evans rat embryos were dissected and disassociated, and primary neurons plated at a density of 6 × 10^5^ cells/mL in 96-well plates, as described previously (Saudou et al., 1998). At *in vitro* day (DIV) 4-5, neurons were transfected with 100 ng of pGW1-mApple (Barmada et al., 2014) to mark cells bodies and 100 ng of an experimental construct (i.e. pGW1-MATR3-EGFP) using Lipofectamine 2000, as before (Barmada et al., 2010). Following transfection, cells were placed into either Neurobasal with B27 supplement (Gibco, Waltham, MA; for all survival experiments) or NEUMO photostable medium (Cell Guidance Systems, Cambridge, UK; for optical pulse labeling experiments). For siRNA knockdown experiments, neurons were transfected with 100 ng of pGW1-mApple per well and siRNA at a final concentration of 90 nM. Cells were treated with either scrambled siRNA (Dharmacon, Lafayette, CO) or siRNA targeting the N-terminal coding region of rat Matr3 (5’ GUC AUU CCA GCA GUC AUC UUU 3’).

### Longitudinal fluorescence microscopy and automated image analysis

Neurons were imaged as described previously (Barmada et al., 2015) using a Nikon (Tokyo, Japan) Eclipse Ti inverted microscope with PerfectFocus3 and a 20X objective lens. Detection was accomplished with an Andor (Belfast, UK) iXon3 897 EMCCD camera or Andor Zyla4.2 (+) sCMOS camera. A Lambda XL Xenon lamp (Sutter) with 5 mm liquid light guide (Sutter Instrument, Novato, CA) was used to illuminate samples, and custom scripts written in Beanshell for use in μManager controlled all stage movements, shutters, and filters. Custom ImageJ/Fiji macros and Python scripts were used to identify neurons and draw regions of interest (ROIs) based upon size, morphology, and fluorescence intensity. Criteria for marking cell death involved rounding of the soma, loss of fluorescence and degeneration of neuritic processes. Custom scripts were also used to identify and draw bounding ROIs around nuclei of transfected cells based upon MATR3-EGFP or Hoechst 33258 (ThermoFisher, Waltham, MA) fluorescence. Coefficient of variation (CV) was calculated as the standard deviation of fluorescence intensity divided by the mean fluorescence intensity within an ROI.

### Immunocytochemistry

Neurons were fixed with 4% paraformaldehyde, rinsed with phosphate buffered saline (PBS), and permeabilized with 0.1% Triton X-100 in PBS. After brief treatment with 10 mM glycine in PBS, cells were placed in blocking solution (0.1% Triton X-100, 2% fetal calf serum, and 3% bovine serum albumin (BSA), all in PBS) at room temperature (RT) for 1 h before incubation in primary antibody, rabbit anti-MATR3 (Abcam EPR10634(B), Cambridge, UK) diluted 1:1000 in blocking solution, overnight at 4 °C. Cells were then washed 3x in PBS and incubated at RT with secondary antibody, goat anti-rabbit 647 (ThermoFisher A-21245) diluted 1:1000 in blocking solution, for 1 h at RT. Following 3x rinses in PBS containing 1:5000 Hoechst 33258 dye (ThermoFisher), neurons were imaged by fluorescence microscopy, as described above.

### Fluorescence recovery after photobleaching

Primary neurons were dissected as above and plated in 8-well borosilicate chambers (LAB-TEK). On DIV 3, they were transfected as before but using 200 μg of pGW1-mApple and 200 μg of pGW1-MATR3-EGFP variants per well. Cell were imaged 2-4 days after transfection using a Nikon A1 confocal microscope operated by Nikon Elements, a 60X objective lens, and a heating chamber with CO_2_ pre-warmed to 37 °C. For MATR3(dRRM1/2)-EGFP variants, an ROI corresponding to half of the granule was outlined with Elements and photobleached using a 488 nm laser set at 30% power, 1 pulse per sec x 7 sec. Fluorescence recovery was monitored up to 10 min after photobleaching. For full-length MATR3 variants, ROIs for photobleaching were drawn in the center of the nucleus for each cell, and recovery was monitored for 6 min.

Image analysis was conducted in FIJI. Rigid body stack registration was used to fix the granules in place relative to the frame. The GFP integrated density for the whole granule was calculated from pre-bleach measurements, as was the fraction of granule integrated density corresponding to the ROI to be photobleached. The decline in this fraction immediately after photobleaching was then calculated and used as the floor, and the return was plotted as the percent recovery within the ROI as a fraction of the original pre-bleach granule integrated density.

Recovery data were fit to the equation y(t) = A(1-e^−Τt^), where A is the return curve plateau, Τ is the time constant, and t is the time post-bleach. The fitted Τ from each curve was then used to calculate the time to half-return (t_1/2_) using the equation t_1/2_ = ln(0.5)/-Τ. To estimate the diffusion coefficient (D) of these molecules, we used the equation D = (0.88w^2^)/(4t_1/2_), where w is the ROI radius (Gopal et al., 2017). This equation assumes spot bleach with a circular stimulation ROI and diffusion limited to the x-y plane. Since we could not be confident that these assumptions were met, we estimated D and downstream parameters by dividing ROI areas by π to approximate w^2^ and solving for D. This estimated value was used in the Einstein-Stokes equation, D = k_B_T/(6πηr), where k_B_ is the Boltzmann constant, T is temperature in K, η is viscosity, and r is the Stokes radius of the particle. As there is no applicable structural data on MATR3, we estimated a Stokes radius of 3.13 nm by applying the MATR3(dRRM1/2)-EGFP fusion protein’s combined molecular weight of 106.4 kDa to the equation R_min_ = 0.66M^1/3^, where R_min_ is the minimal radius in nm of a sphere that could bound a globular protein with a molecular weight of M (Erickson, 2009). Using these constants and the estimated D for each granule, the Einstein-Stokes equation was rearranged to solve for η.

Photobleaching data from full-length MATR3-EGFP was analyzed in a similar fashion. After calculating the nuclear integrated density, the fraction attributable to photobleaching within the ROI was used for normalization. Intensity data were fit to the y(t) = A(1-e^−⊤t^) equation, t_1/2_ values were calculated as before, and D determined by the equation D = (0.88w^2^)/(4t_1/2_).

### Nuclear/cytoplasmic fractionation and differential solubility

HEK293T cells were transfected in a 6-well plate with 3 μg of DNA per well using Lipofectamine 2000 according to the manufacturer’s instructions. For nuclear/cytoplasmic fractionation, cells were washed with cold PBS 24 h after transfection, collected with resuspension buffer (10 mM Tris, 10 mM NaCl, 3 mM MgCl_2_, pH 7.4), and transferred to a pre-chilled 1.5 mL conical tube to sit on ice for 5 min. An equal volume of resuspension buffer with 0.6% Igepal (Sigma, St. Louis, MO) was then added to rupture cell membranes and release cytoplasmic contents, with occasional inversion for 5 min on ice. Nuclei were pelleted at 100 × g at 4 °C for 10 min using a tabletop centrifuge. The supernatant (cytosolic fraction) was collected, and the nuclei were rinsed twice in resuspension buffer without Igepal. To collect nuclear fractions, pelleted nuclei were lysed in RIPA buffer (Pierce) with protease inhibitors (Roche, Mannheim, Germany) on ice for 30 min with occasional inversion. Samples were centrifuged at 9,400 × g at 4 °C for 10 min, and the supernatant was saved as the nuclear fraction.

For differential solubility experiments, transfected HEK293T were collected in cold PBS 24 h after transfection and transferred to a pre-chilled conical tube on ice. Cells were then centrifuged at 100 × g for 5 min at 4 °C to pellet cells, the PBS was aspirated, and cells were resuspended in RIPA buffer with protease inhibitors. Following lysis on ice for 15 min with occasional inversion, cells were sonicated at 80% amplitude with 5 sec on/5 sec off for 2 min using a Fisherbrand Model 505 Sonic Dismembrenator (ThermoFisher). Samples were centrifuged at 41,415 × g for 15 min at 4 °C to pellet RIPA-insoluble material, with the supernatant removed and saved as the RIPA-soluble fraction. The RIPA-insoluble pellet was washed in RIPA once, and contents resuspended vigorously in urea buffer (7 M urea, 2 M thiourea, 4% CHAPS, 30 mM Tris, pH 8.5). Samples were again centrifuged at 41,415 × g for 15 min at 4 °C, and the supernatant was saved as the RIPA-insoluble, urea-soluble fraction.

For SDS-PAGE, stock sample buffer (10% SDS, 20% glycerol, 0.0025% bromophenol blue, 100 mM EDTA, 1 M DTT, 20 mM T ris, pH 8) was diluted 1:10 in lysates and all samples except urea fractions were boiled for 10 min before 5-15 μg of protein were loaded onto 4-15% gradient gels (Bio-Rad, Hercules, CA). For urea fractions, total protein concentration was too low to quantify and so equal volumes of sample across conditions were mixed 1:1 with water and loaded. After electrophoresis, samples were transferred at 30 V overnight at 4 °C onto an activated 2 μm nitrocellulose membrane (Bio-Rad), blocked with 3% BSA in 0.2% Tween-20 in Tris-buffered saline (TBST), and blotted overnight at 4 °C with the following primary antibodies: rabbit anti-MATR3 (Abcam EPR10634(B)), mouse anti-GAPDH (Millipore Sigma MAB374), and rabbit anti-H2B (Novus NB100-56347), all diluted 1:1000 in 3% BSA, 0.2% TBST. The following day, blots were washed in 0.2% TBST, incubated at RT for 1 h with AlexaFluor goat anti-mouse 594 (ThermoFisher A-11005) and goat anti-rabbit 488 (ThermoFisher A-11008), both diluted 1:10,000 in 3% BSA in 0.2% TBST. Following treatment with secondary antibody, blots were washed in 0.2% TBST, placed in Tris-buffered saline, and imaged using an Odyssey CLx Imaging System (LI-COR, Lincoln, NE).

### Statistical analysis

Statistical analyses were performed in R or Prism 7 (GraphPad). For primary neuron survival analysis, the publically available R survival package was used to determine hazard ratios describing the relative survival among populations through Cox proportional hazards analysis. For half-life calculations, a custom R script was applied to fit log-transformed TRITC intensity data to a linear equation. Photobleaching recovery data were fit to the y(t) = A(1-e^−⊤t^) equation using non-linear regression in R. siRNA knockdown data were plotted using Prism 7, and significance determined via the twotailed t-test. One-way ANOVA with Tukey’s post-test was used to assess for significant differences among nuclear/cytoplasmic ratios, viscosities, D values, and half-lives. Data are shown as mean ± SEM unless otherwise stated.

**Table 1.**
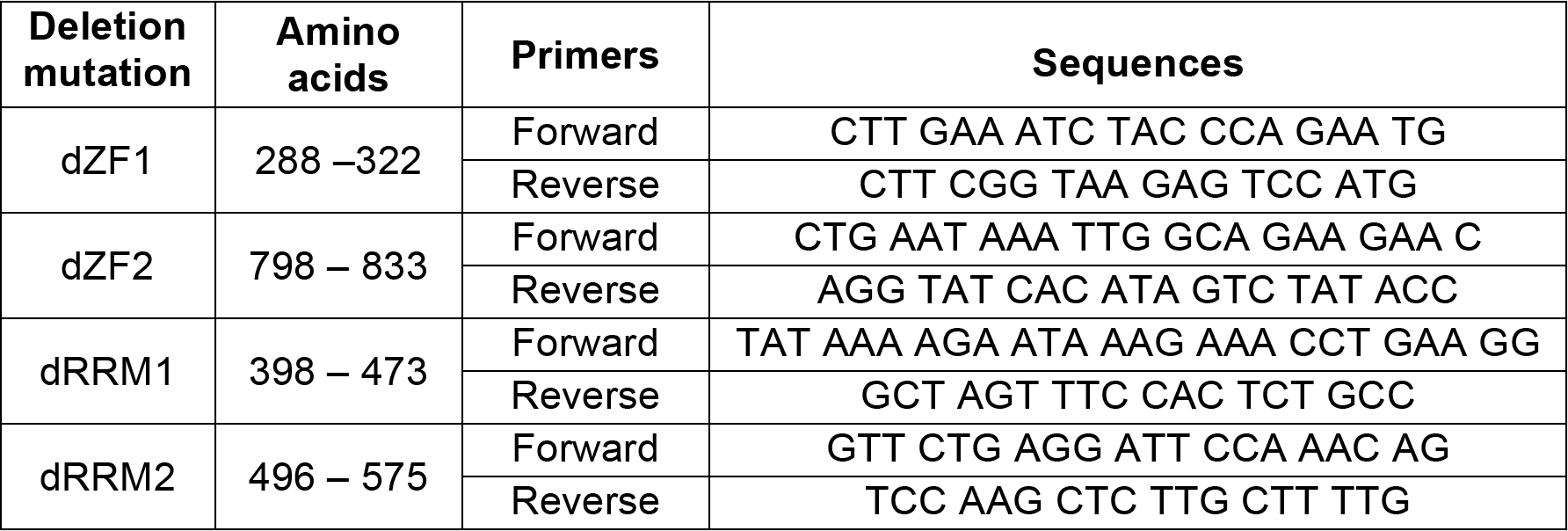

**Table 2.**
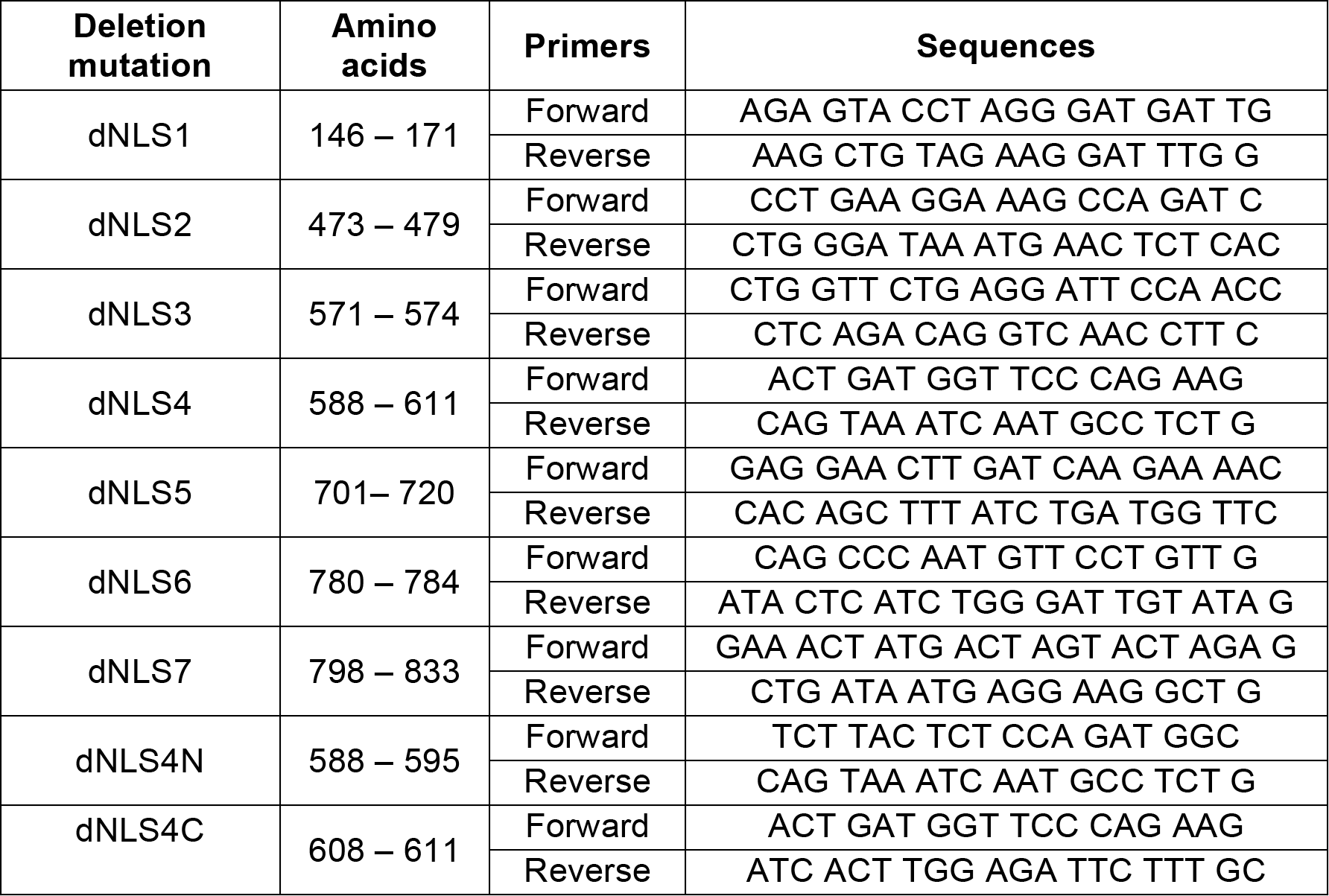

**Table 3.**
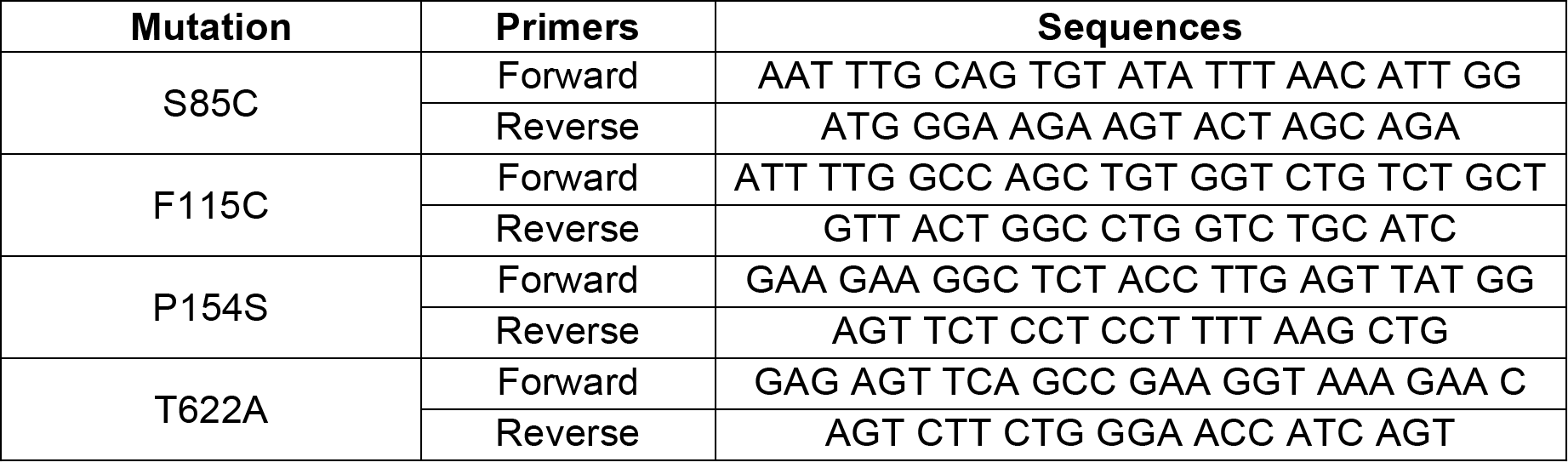

## ACKNOWLEDGEMENTS

We thank Dr. Stephen Lentz for his assistance with confocal microscopy, Drs. Claudia Figueroa-Romero and Hilary Archbold for their experimental advice, and Brittany Flores for technical assistance in assembling the manuscript.

The present work was supported in part by funding from the National Institutes of Health (NIH) National Institute for Neurological Disorders and Stroke (NINDS) R01 NS097542 (S.J.B.), National Institute for Aging (NIA) P30 AG053760 (S.J.B.), and National Institute of General Medical Sciences (NIGMS) T32 GM007863 (A.M.M.) and T32 NS076401 (A.M.M.); the Protein Folding Diseases Initiative at the University of Michigan (S.J.B.); the Program for Neurology Research and Discovery (E.L.F.); and the A. Alfred Taubman Medical Research Institute (S.J.B., E.L.F.). Confocal microscopy was performed at the Microscopy & Image Analysis Core of the Michigan Diabetes Research Center funded by NIH grant P60DK020572 from the National Institute of Diabetes and Digestive and Kidney Diseases (NIDDK).

## AUTHOR CONTRIBUTIONS

A.M.M., Y.S.H., E.L.F., and S.J.B. designed the study; R.A.M. wrote original code for data analysis; X.L. performed primary neuron isolations; Y.S.H. created all MATR3 constructs and identified NLS-like sequences within MATR3; A.M.M. and Y.S.H. conducted neuronal survival experiments; A.M.M. performed all confocal microscopy and data analysis, and assembled all figures; A.M.M. and S.J.B. wrote the manuscript; A.M.M., S.J.B., Y.S.H and E.L.F. edited and revised the manuscript.

